# Inhibitory effect of the TSG-6 on the BMP-4/Smad signaling pathway and odonto/osteogenic differentiation of dental pulp stem cells

**DOI:** 10.1101/2020.03.17.995274

**Authors:** Ying Wang, Shuai Yuan, Jingjing Sun, Yuping Gong, Sirui Liu, Runying Guo, Wei He, Yiming Liu, Peng Kang, Rui Li

**Affiliations:** Department of Stomatology, The First Affiliated Hospital of Zhengzhou University, Zhengzhou University

**Keywords:** BMP/Smad, dental pulp stem cells, pulpitis, tumor necrosis factor-inducible protein 6

## Abstract

This study aimed to observe the molecular mechanism underlying the effect of tumor necrosis factor–inducible protein 6 (TSG-6) on the bone morphogenetic protein-4 (BMP-4)/drosophila mothers against decapentaplegic protein(Smad) signaling pathway and mineralization of dental pulp stem cells (DPSCs) in inflammatory environment. Normal and TSG-6 gene–modified DPSCs were cultured in a mineralization-inducing fluid containing 0 and 50 ng/mL TNF-α separately. The real-time polymerase chain reaction was used to measure the expression of TSG-6 and odonto/osteogenic differentiation makers at the mRNA level. Western blot analysis and cellular immunofluorescence were used to observe the odonto/osteogenic differentiation of DPSCs and the variation of BMP-4/Smad signaling pathway at the protein level. Moreover, normal and modified DPSCs combined with hydrogel were used for subcutaneous implantation in nude mice. The expression of odonto/osteogenic markers and BMP-4/Smad-related proteins was lower in Ad-TSG-6 DPSCs than in normal DPSCs after mineralization induction, and was higher in TSG-6-RNAi DPSCs than in normal DPSCs after culturing with mineralization-inducing fluid containing 50 ng/mL TNF-α. The subcutaneous transplantation of normal and modified DPSCs combined with hydrogel in nude mice demonstrated that normal DPSCs were formed in the tissue containing collagen. The tissue formed by Ad-TSG-6 DPSCs was highly variable, and the cells were very dense. The expression of odonto/osteogenic markers of Ad-TSG-6 DPSCs were lower in Ad-TSG-6 DPSCs than in normal DPSCs. We can know that TNF-α regulates the expression of TSG-6, thereby inhibiting the BMP-4/Smad signaling pathway and the odonto/osteogenic differentiation ability of DPSCs.

## Introduction

Pulpitis is an acute or chronic inflammation of the teeth caused by caries or trauma(1).(2) It can develop into periapical inflammation and affect periodontal health, ultimately leading to tooth loss. The incidence of pulpitis continues to increase every year; it has become a common disease endangering oral and even human health (3). The World Health Organization has listed pulpitis as a key disease to be prevented and treated. Root canal treatment is one of the most common treatments for pulpitis(4,5). However, this method can only remove the diseased pulp(6). Hence, developing alternative treatment options for pulpitis treatment is a critical medical need.

Dental tissue engineering is an emerging field that considers dental pulp regeneration as a new solution for dental problems(7-9). A previous study examined the regeneration of dental pulp *in vivo(10,11)*, indicating that dental pulp stem cells (DPSCs) had the potential for treating pulp disease. However, the existing findings rarely target the repairing ability and mechanism of DPSCs under inflammatory conditions. At present, most scholars believe that the number of cells differentiated from mesenchymal stem cells can be increased in a weak inflammatory environment, while their ability to differentiate is weakened in a strong inflammatory environment(11-13). Moreover, some evidence indicates that the odonto/osteogenic differentiation of DPSCs is affected by the extracellular microenvironment(14). However, the mechanism has not been fully understood yet.

Tumor necrosis factor-alpha (TNF-α) is one of the most important factors involved in the process of inflammation(15,16). Moreover, it inhibits the osteogenic differentiation of mesenchymal stem cells at a high concentration(17,18). It has been proved to inhibit osteogenesis in a number of previous studies(17,19). Therefore, it was used to simulate the inflammatory microenvironment in the present study. Tumor necrosis factor-inducible protein 6 (TSG-6) is induced mainly by TNF-α and interleukin 1, and is an important anti-inflammatory factor(20). TSG-6 is a cytokine produced by mesenchymal stem cells. It plays an important role against inflammation and in disease treatment(21). It can reverse liver fibrosis and ameliorate liver damage in mice. Furthermore, TSG-6 plays a role in the treatment of autoimmune diseases(22,23). Despite its potent anti-inflammatory properties, TSG-6 has a reverse effect on the differentiation of mesenchymal stem cells(24). Studies have shown that mesenchymal stem cells (MSCs) expressing higher levels of TSG-6 have a lower osteogenic capacity(25,26). During the progression of pulpitis, more destruction occurs than repair. Improving the mineralization ability of dental pulp stem cells in inflammatory environment could promote tne repair process. Hence, the effect of high expression of TSG-6 caused by strong inflammation on the dental/osteogenic differentiation ability of DPSCs and its mechanism should be explored.

Bone morphogenetic protein (BMP)/Smad signaling pathway is an important molecular mechanism for the odonto/osteogenic differentiation of DPSCs. BMP-4 is one of the important molecules affecting early odontogenic differentiation(27). TSG-6 can inhibit bone morphogenetic protein-2 (BMP-2)-mediated osteoblast differentiation(28). However, the effect of TSG-6 on the differentiation of odontogenic stem cells has not been studied, and whether TSG-6 can affect the BMP-4/Smad signaling pathway in DPSCs is still unknown. In addition, there were few studies on how to improve the mineralization ability of DPSCs in inflammatory environment. The purpose of this study was to observe the molecular mechanism underlying the effect of TSG-6 on the mineralization of DPSCs. The results might provide a new idea for the treatment of pulpitis.

## Results

### Isolation, culture, and identification of DPSCs

Cellular immunofluorescence showed that DPSCs expressed Vimentin and STRO-1 proteins (Fig. 1Aa and 1Ab), which were specific proteins expressed in MSCs. However, the obtained cells were negative for the epithelial cell marker CK-14 (Fig. 1Ac). After osteogenic induction, the cells produced mineralized nodules (Fig. 1Ad) and lipid droplets after adipogenic induction (Fig. 1Ae), indicating that the obtained cells had multidirectional differentiation potential. Moreover, the flow cytometry results showed that the obtained cells expressed MSC markers, including CD73 and CD90, but they did not express endothelial cell markers, including CD31, CD34, and T cell marker CD3 (Fig. 1B). These findings indicated that the isolated and cultured cells were DPSCs.

**Figure 1.**
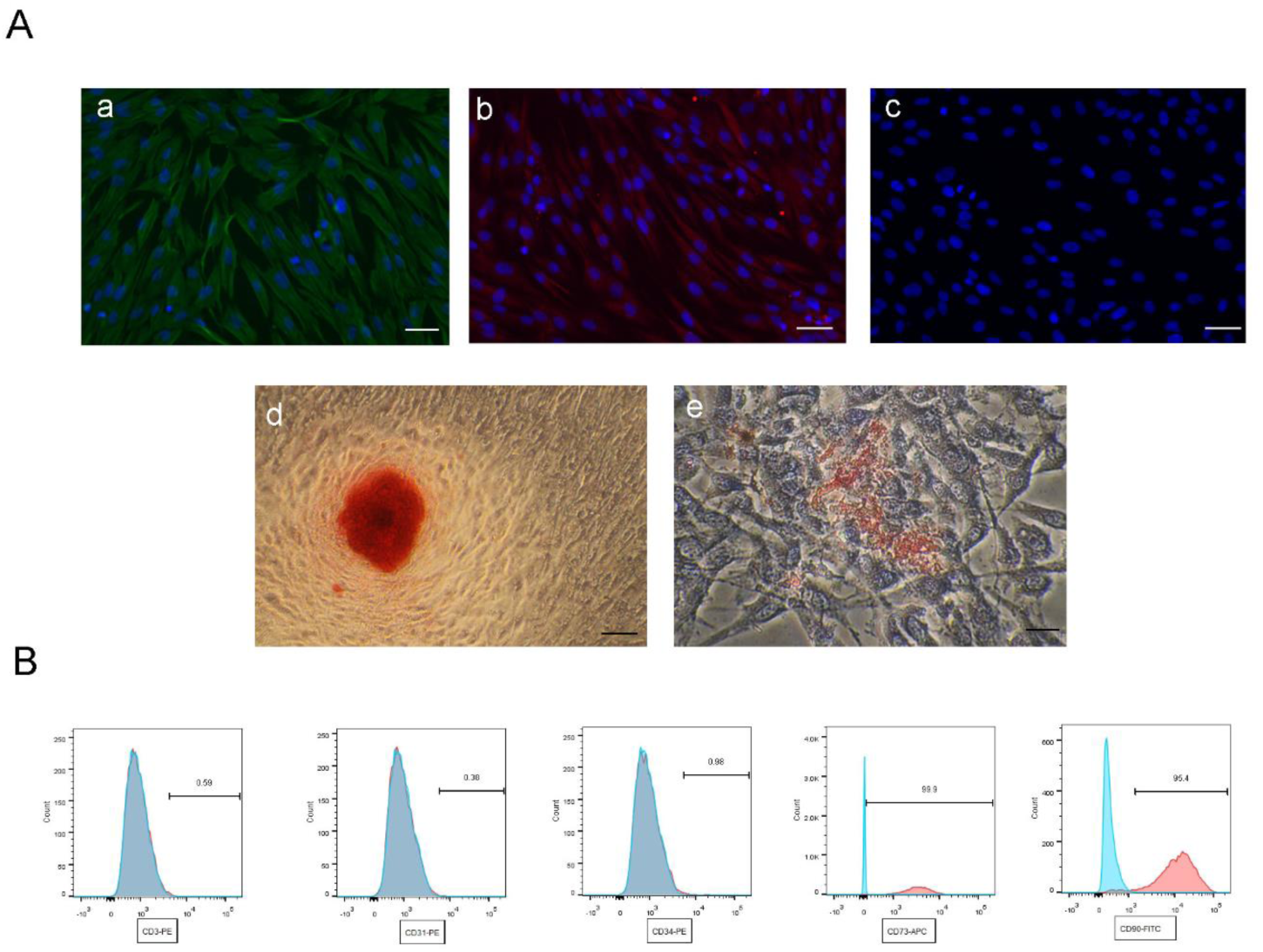
Cells expressed Vimentin (Aa) and STRO-1 (Ab) proteins but did not express the epithelial cell marker CK-14 (Ac). They showed (Ad) mineralized nodules (Ae) when they were cultured in osteogenic and adipogenic induction medium for 21 and 18 days. Flow cytometry (B) indicated that the cells were positive for mesenchymal stem cell surface markers CD73 and CD90 but negative for endothelial cell markers CD31 and CD34 and T cell marker CD3. Scale bars of Aa, Ab, Ac, and Ad = 40 μm. Scale bars of Ae =20 μm.

### Inflammatory DPSCs highly expressed TSG-6

The inflammatory DPSCs could highly express the anti-inflammatory factor TSG-6 (Fig. 2Aa), indicating that TSG-6 played an important role in the inflammatory environment. Alkaline phosphatase is an important indicator of the odonto/osteogenesis differentiation of DPSCs. The phosphatase activity of DPSCs significantly reduced in the mineralization-inducing fluid containing 50 ng/mL TNF-α solution (Fig. 2Ab). Therefore, 50 ng/mL solution was used as the concentration induced *in vitro* by TNF-α. The treatment of DPSCs with 50 ng/mL TNF-α solution reduced the expression levels of odonto/osteogenesis-related genes (Fig. 2Ac), indicating that high concentrations of TNF-α could reduce the odonto/osteogenic ability of DPSCs.

**Figure 2.**
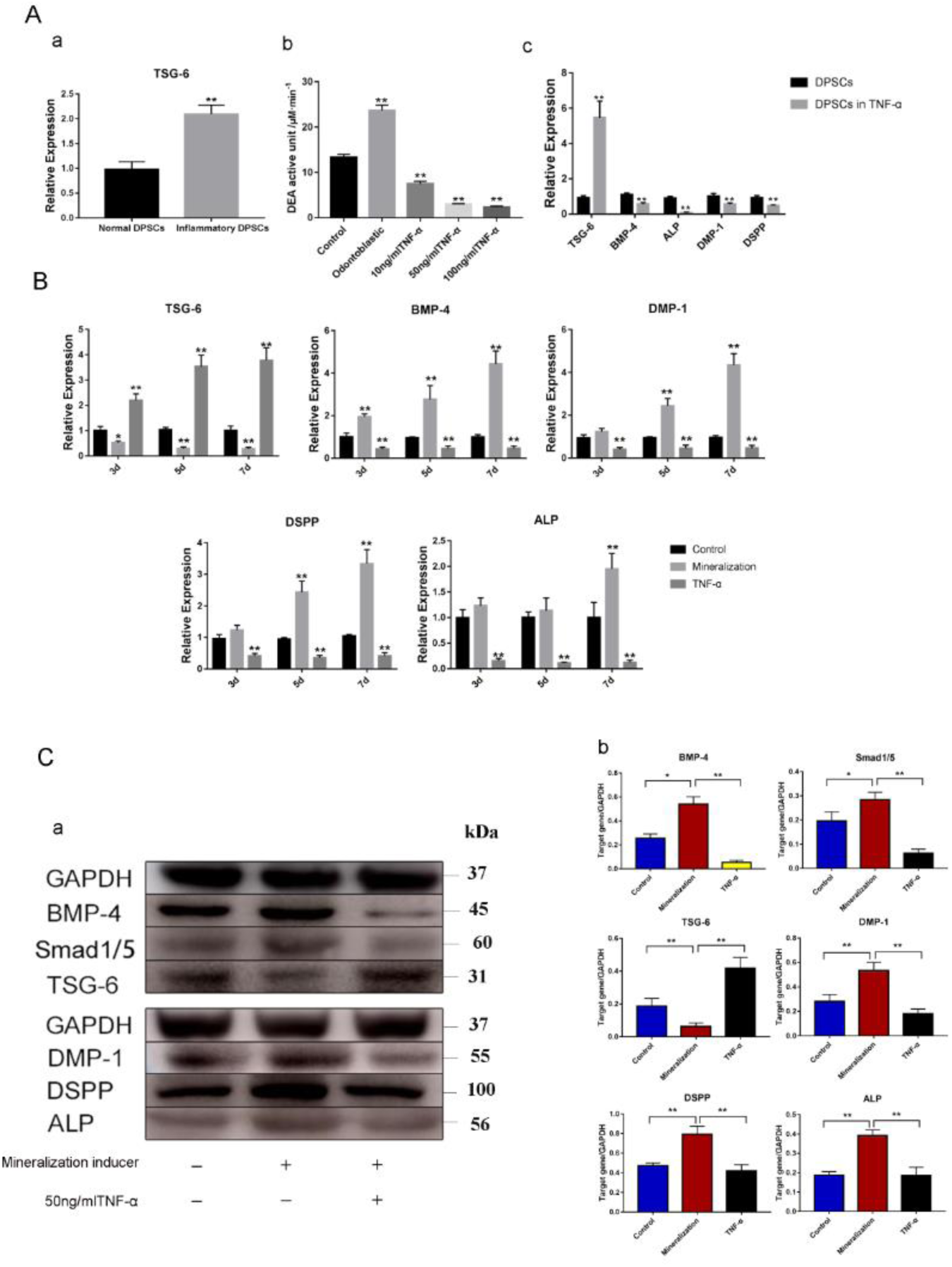
Inflammatory dental pulp stem cells (DPSCs) could highly express TSG-6 (Aa). In mineralization-inducing fluid containing TNF-α solution, the phosphatase activity of DPSCs significantly reduced (Ab). Treatment of DPSCs with 50 ng/mL TNF-α reduced the expression of odonto/osteogenesis-related genes (Ac). Real-time PCR (B) was performed to assess the effect of the 50 ng/mL TNF-α on the odonto/osteogenesis of DPSCs after 3, 5, and 7 days, respectively. BMP-4, DMP-1, DSPP, and ALP were upregulated to different degrees in cells induced by the mineralization inducer and downregulated to varying degrees in cells induced by the mineralization-inducing fluid containing 50 ng/mL TNF-α. Western blot analysis (Ca) was performed after 48 h and 14 days, and the relative levels were quantified (Cb). BMP-4, phosphorylated Smad1/5, DMP-1, DSPP, and ALP were upregulated, while the expression levels of TSG-6 decreased, after treatment with mineralization-inducing fluid, contrary to the changes in the levels in DPSCs treated with mineralization-inducing fluid containing 50 ng/mL TNF-α (*P < 0.05, **P < 0.01, compared with normal DPSCs without induction).

### High concentrations of TNF-α could increase the expression level of TSG-6 and decrease the expression level of odonto/osteogenesis markers of DPSCs

The expression levels of BMP-4, DMP-1, DSPP, and ALP showed that the mineralization-induced dentate-related genes were upregulated to different degrees (Fig. 2B). However, the odonto/osteogenesis-related genes were downregulated to varying degrees induced by the mineralization-inducing fluid containing 50ng/mL TNF-α, indicating that high concentrations of TNF-α could inhibit the odonto/osteogenic differentiation of DPSCs. Moreover, the expression of TSG-6 and signaling pathway-related proteins BMP-4 and phosphorylated Smad1/5 was detected by Western blot analysis after 48 h of induction. The expression of odonto/osteogenesis-related proteins, including DMP-1, DSPP, and ALP, was detected after 14 days of induction. After 48 h of odonto/osteogenesis induction, the expression level of TSG-6 protein decreased, while the expression level of BMP-4 increased (Fig. 2Ca and 2Cb). Moreover, the phosphorylation level of Smad1/5 protein increased. However, the expression level of TSG-6 increased in the group induced with TNF-α, while the expression level of BMP-4 and the phosphorylation of Smad l/5 protein decreased. The results indicated that the expression level of TSG-6 in DPSCs decreased during the process of odonto/osteogenesis. Therefore, TSG-6 might serve as a negative regulatory protein in the differentiation of DPSCs. Furthermore, high concentrations of TNF-α could inhibit the BMP-4/Smad signaling pathway. After 14 days of odonto/osteogenesis induction, the expression of odonto/osteoblast-associated proteins increased to variable degrees. The results indicated that high concentrations of TNF-α could inhibit the odonto/osteogenesis ability of DPSCs.

### TSG-6 could inhibit the BMP-4/Smad signaling pathway and odontogenic/osteogenic differentiation in DPSCs

The lentivirus had no effect on the results of this study; the data on the effect of lentivirus transfection are provided in Supplementary Material. The expression level of BMP-4 protein and the phosphorylation level of Smad1/5 were lower in Ad-TSG-6 DPSCs than in normal DPSCs after mineralization induction (Fig. 3), indicating that TSG-6 could inhibit the BMP-4/Smad signaling pathway. However, the expression level of BMP-4 protein and the phosphorylation level of Smad1/5 increased in TSG-6-RNAi DPSCs compared with normal DPSCs after induction with mineralization-inducing fluid containing 50 ng/mL TNF-α. The expression of DMP-1, DSPP, and ALP in each component was analyzed using Western blot analysis. The expression of odonto/osteogenesis-related proteins decreased in Ad-TSG-6 DPSCs compared with normal DPSCs after induction with mineralization-inducing fluid (Fig. 3), indicating that TSG-6 could inhibit the odonto/osteogenic differentiation of DPSCs. Moreover, the expression level of odonto/osteogenesis-related proteins increased to varying degrees in TSG-6-RNAi DPSCs compared with normal DPSCs after induction with the mineralization-inducing fluid containing 50 ng/mL TNF-α. The results indicated that the high concentrations of TNF-α could inhibit the BMP-4/Smad signaling pathway and the odonto/osteogenic differentiation of DPSCs by promoting the high expression of TSG-6. The immunofluorescence showed that the expression levels of DMP-1 and DSPP were lower in Ad-TSG-6 DPSCs than in normal DPSCs, while the levels were higher in TSG-6-RNAi DPSCs than in normal DPSCs (Figure 4).

**Figure 3.**
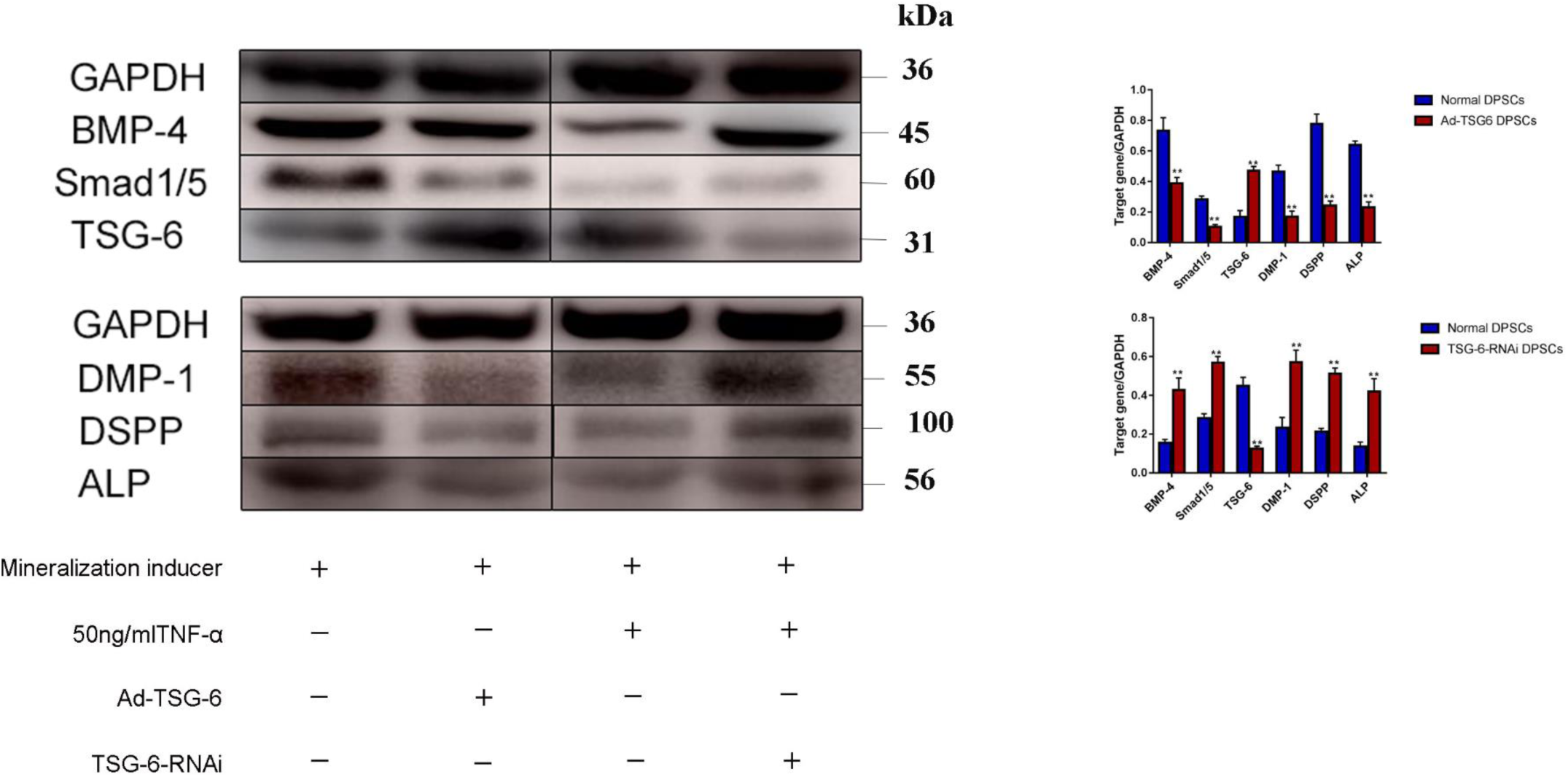
Western blot analysis was performed after 48 h and 14 days, and the relative levels were quantified. The expression levels of BMP-4 and phosphorylated Smad1/5 were lower in Ad-TSG-6 DPSCs than in normal DPSCs after mineralization induction; the expression levels of DMP-1, DSPP, and ALP were also lower. This was contrary to the levels in TSG-6-RNAi DPSCs treated with mineralization-inducing fluid containing 50 ng/mL TNF-α (*P < 0.05, **P < 0.01, compared with normal DPSCs with induction). Scale bars =200 μm.

**Figure 4.**
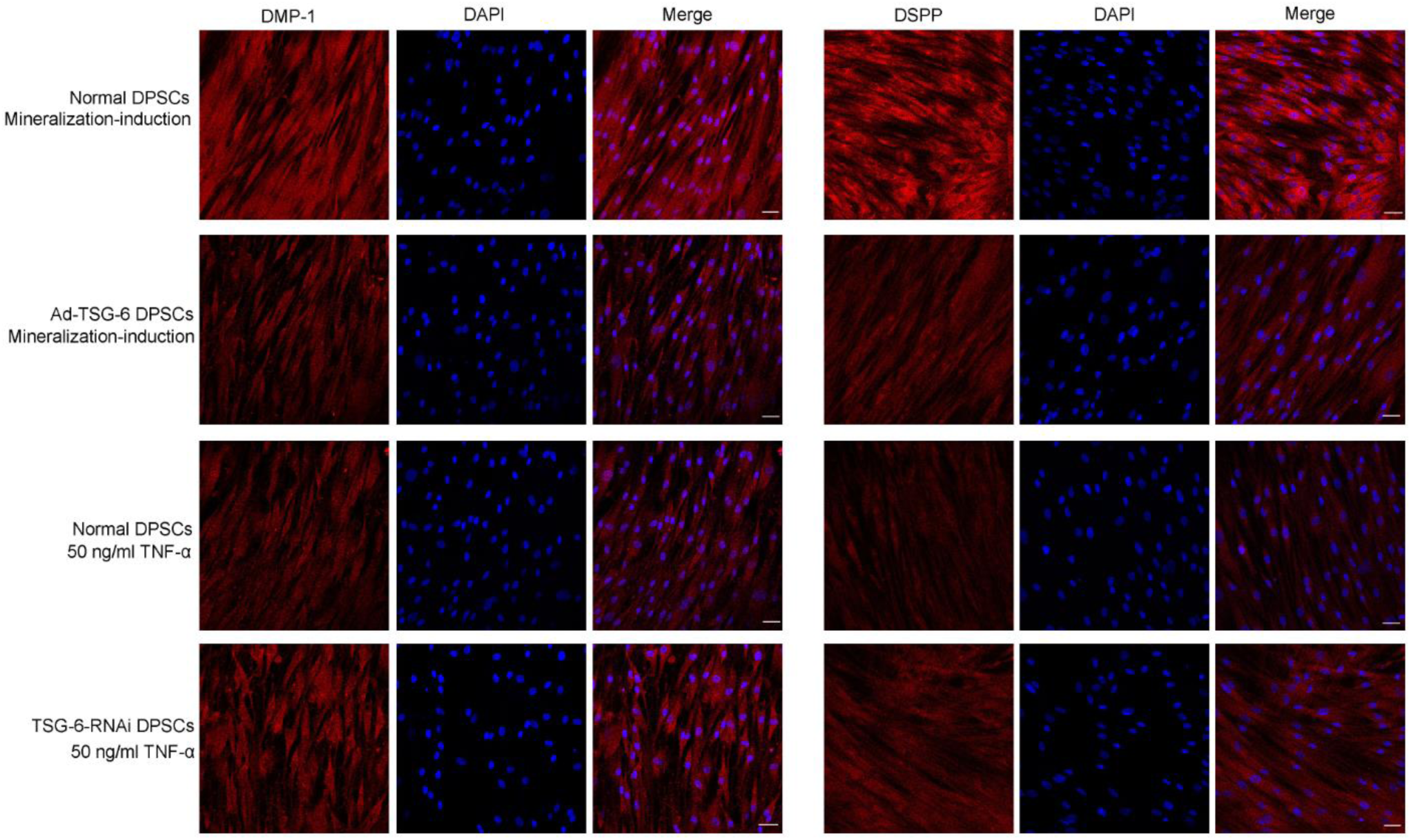
Immunofluorescence was performed to analyze the expression of DMP-1 and DSPP in each group after 14 days of induction. The expression levels were lower in Ad-TSG-6 DPSCs than in normal DPSCs after mineralization induction, and while the levels were higher in TSG-6-RNAi DPSCs than in normal DPSCs (P < 0.05). Scale bars =40 μm.

### Subcutaneous ectopic osteogenesis in nude mice

DPSCs combined with hydrogel mixture were implanted into the dorsum of the nude mice, and the grafts were collected after 6 weeks. However, only osteogenesis-induced normal DPSCs and Ad-TSG-6 DPSCs produced ectopic tissue (Fig. 5A). HE and Masson staining were used to observe the morphology and structure of subcutaneous grafts. After induction with the mineralization-inducing fluid, normal DPSCs were formed in tissue containing collagen (Fig. 5Ba and 5Bb). The tissue formed by Ad-TSG-6 DPSCs was highly variable, and the cells were very dense (Fig. 5Bc and 5Bd). However, normal and TSG-6-RNAi DPSCs induced by mineralization-inducing fluid containing 50 ng/mL TNF-α did not form ectopic tissue subcutaneously in the nude mice. Only muscle tissues were observed (Fig. 5Be–5Bh), indicating that this was significantly associated with the inhibition by high concentrations of TNF-α. Moreover, the immunohistochemical staining was used to evaluate the expression of odonto/osteogenic-related genes, including OCN, OPN, collagen I, DMP-1, collagen II, BSP, RUNX2, and DSPP. Normal DPSCs induced by mineralization fluid had high expression levels of osteogenesis-related indicators (Fig. 6). The positive expression was significantly lower in Ad-TSG-6 DPSCs than in normal DPSCs.

**Figure 5.**
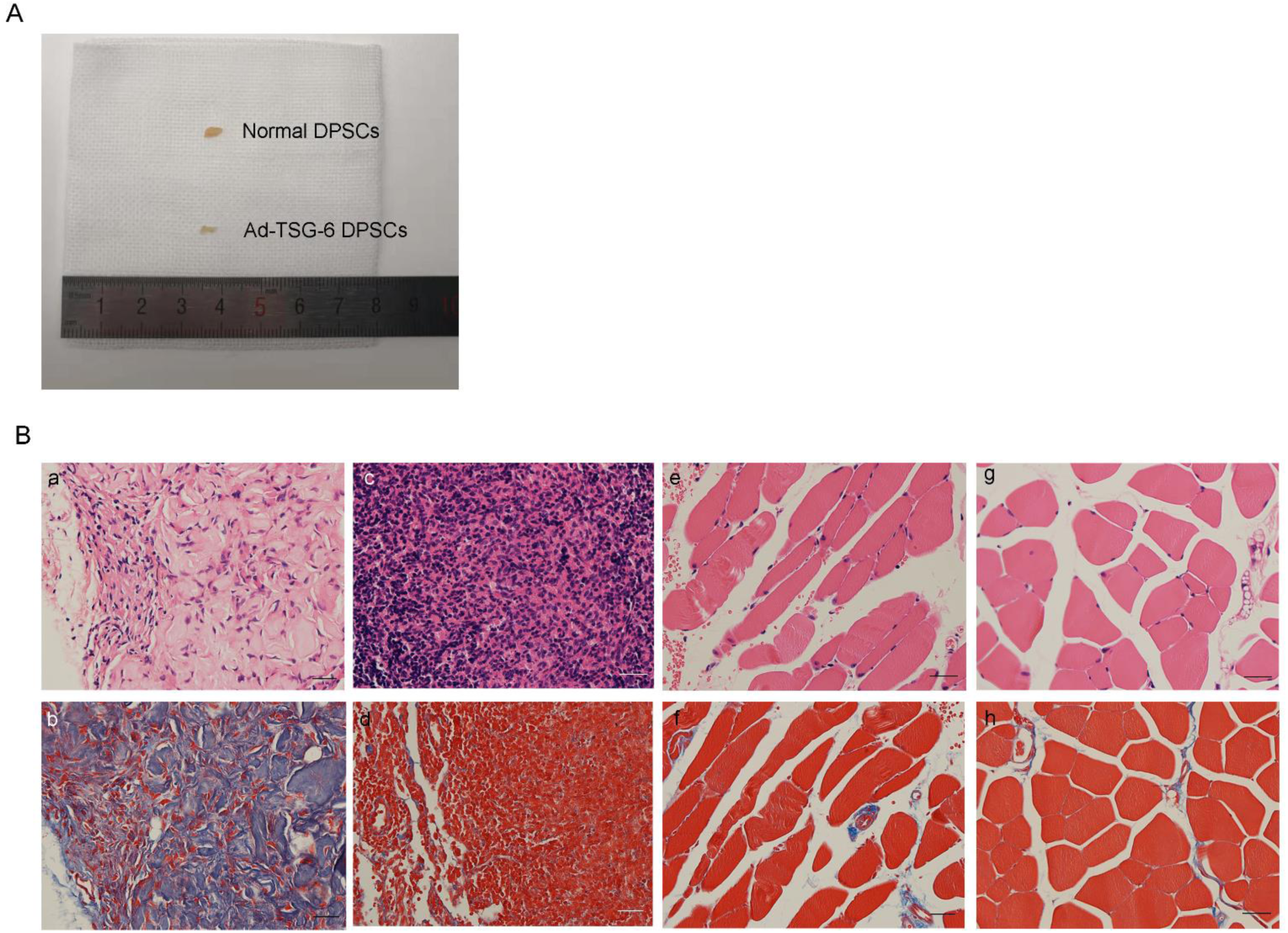
Grafts were collected after 6 weeks (A). HE (Ba, Bc, Be, and Bg) and Masson (Bb, Bd, Bf, and Bh) staining showed that normal DPSCs were formed in the tissue containing abundant collagen, while the tissues formed by Ad-TSG-6 DPSCs were highly variable and the cells were very dense. Scale bars = 20 μm.

**Figure 6.**
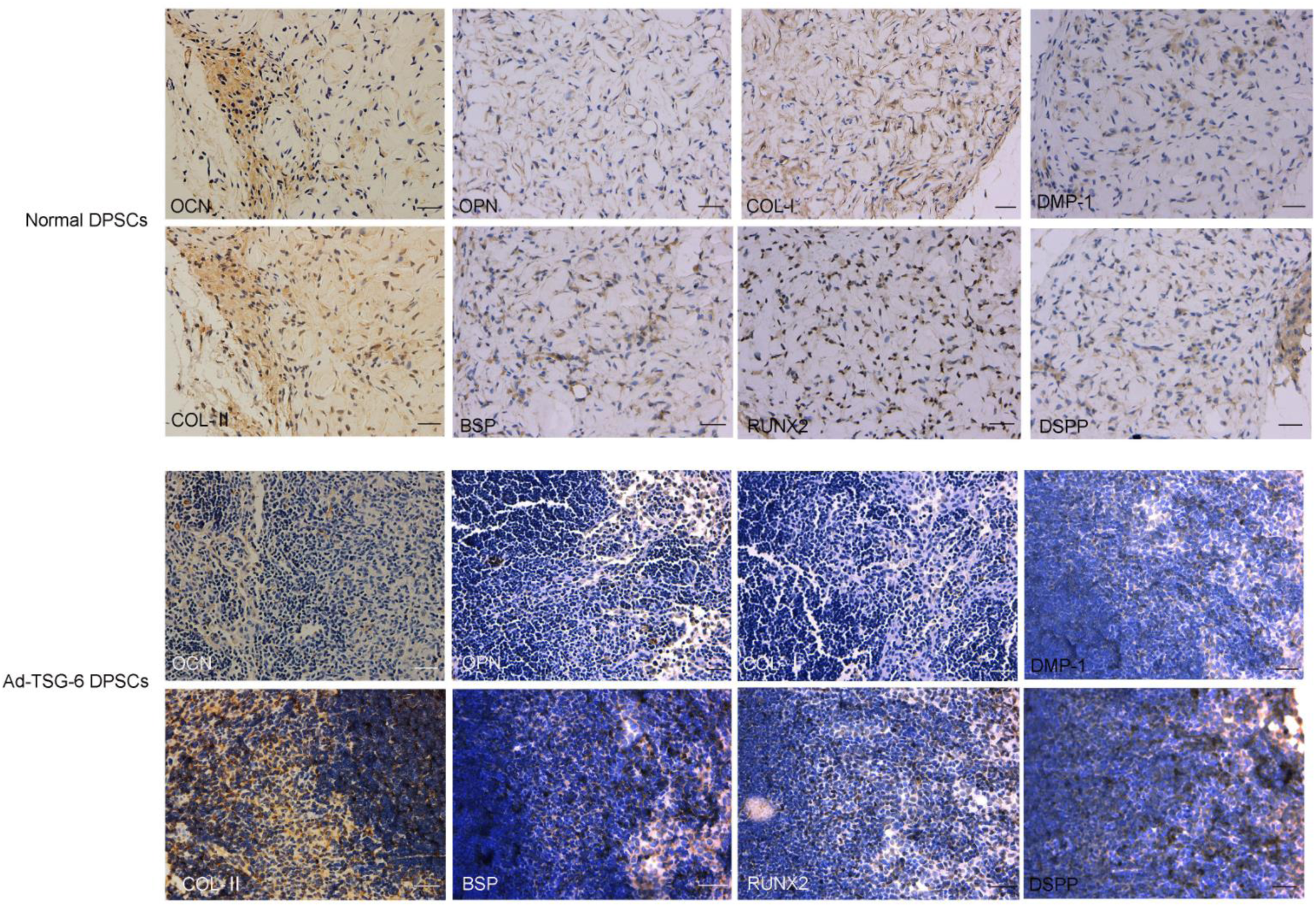
Immunohistochemical evaluation of the subcutaneous ectopic osteogenesis in nude mice was performed after 8 weeks. The expression levels of OCN, OPN, collagen I, DMP-1, collagen II, BSP, RUNX2, and DSPP were lower in Ad-TSG-6 DPSCs than in normal DPSCs (P < 0.05). Scale bars = 20 μm.

## Discussion

In recent years, scholars have increasingly focused on the potential of mesenchymal stem cells in disease treatment(29,30). These cells can be used to treat cardiovascular diseases, tumors, and autoimmune diseases(31-34). As a source of mesenchymal stem cells, DPSCs also have significant potential in therapeutics and tissue regeneration(35-37). Scholars have used the exfoliated deciduous DPSC transplantation to treat injured immature permanent teeth and complete the pulp regeneration process, including the odontoblastic layer, nerves, and blood vessel formation(10). The DPSCs can be used in studies on pulp diseases(38,39). However, the extracellular microenvironment, especially the inflammatory microenvironment, can affect the differentiation ability of mesenchymal stem cells(40,41). Therefore, studying the changes in DPSCs in the inflammatory microenvironment and the underlying molecular mechanism is of great significance for the treatment of pulp inflammation. The present study considered factors that could inhibit the differentiation ability of DPSCs in an inflammatory environment as the breakthrough point, so as to find a novel method for treating pulp inflammation. The present study demonstrated that the TNF-α regulated the BMP-4/Smad signaling pathway and the odonto/osteogenic differentiation ability of DPSCs by regulating the expression of TSG-6.

BMP-4, a member of the transforming growth factor-beta superfamily, is expressed in the presumptive dental epithelium at the initiation of tooth development(42). BMP signaling plays a significant role in early tooth development, which is evidenced by the disruption of BMP signaling leading to an early arrested tooth development(43). In the present study, in the mineralization-inducer with high concentration of TNF-α, the expression of genes and proteins of BMP-4, ALP, DMP-1 and DSPP were reduced, and BMP-4/Smad signal pathway-related proteins were reduced, while the expression of genes and proteins of TSG-6 were increased. It was demonstrated that the expression of TSG-6 were opposite to that of odonto/osteogenesis-related genes and proteins and BMP-4/Smad pathway. Then we modified TSG-6 gene in DPSCs, and made further researech. Ad-TSG-6 DPSCs showed reduced expression levels of odonto/osteogenesis-related genes and proteins, indicating that TSG-6 reduced the mineralization ability of DPSCs. These results were consistent with the literature reports(28,44). Moreover, the variations in the BMP-4/Smad signaling pathway were detected during this process, and the signaling pathway was inhibited when the expression level of TSG-6 increased. After the TSG-6 gene knock-down, Western blot analysis was performed. TSG-6-RNAi DPSCs showed increased expression levels of BMP-4/Smad signaling pathway- and odonto/osteogenesis-related proteins on treatment with high-concentration TNF-α, indicating that TNF-α inhibited the differentiation of DPSCs, which was a result of the high expression of TSG-6. Then, immunofluorescence was performed. The results showed that the expression levels of DMP-1 and DSPP were lower in Ad-TSG-6 DPSCs than in normal DPSCs after odonto/osteogenesis induction. While the levels were higher in TSG-6-RNAi DPSCs than in normal DPSCs after induced by mineralization-inducer with 50ng/ml TNF-α. In vitro experiment, TSG-6 could inhibit the BMP-4/Smad signaling pathway and mineralization ability of DPSCs. Because BMP-4/Smad signaling pathway is an important regulatory pathway for odontogenic / osteogenic differentiation, TSG-6-induced decreased mineralization of DPSCs may be caused by inhibition of BMP-4/Smad signaling pathway. However, we need further study to make the activation or inhibition of BMP-4/Smad signaling pathway in DPSCs to verify this speculation. In our present study, if the expression level of TSG-6 was decreased, DPSCs could have higher mineralization ability in the stronger inflammatory environment, and this may be considered as a reference to improve the differentiation ability of DPSCs in inflammatory environment and the treatment of pulpitis.

To enhance the reliability of the study, the complexes that combined DPSCs and hydrogel were implanted in nude mice to observe subcutaneous osteogenesis. A hydrogel is a material with good biocompatibility; it can be used for stem cell transplantation and can provide scaffolds for stem cells. The hydrogel has no cytokines and hence had no additional effect in this study. However, only osteogenesis-induced normal DPSCs and Ad-TSG-6 DPSCs produced ectopic tissue, which might be attributed to the high concentrations of TNF-α that could attenuate the odonto/osteogenic differentiation of DPSCs(45). Our vitro experiments can also illustrate that, in the mineralization-inducer with 50ng/ml TNF-α, the odonto/osteogenic ability of’ both nomal DPSCs and TSG-6-RNAi DPSCs was reduced very obviously. Because of the significantly reduced mineralization ability, there were no ectopic tissue in these two groups in vivo experiment. The tissues formed by Ad-TSG-6 DPSCs were highly variable and the cells were very dense, while the tissues formed by normal DPSCs containing abundant collagen. This might be because the high expression of TSG-6 inhibited the mineralization ability of DPSCs to affect the formation of the subcutaneous ectopic bone tissue. The expression of odonto/osteoblast markers, including OCN, OPN, collagen I, DMP-1, collagen II, BSP, RUNX2, and DSPP, were determined by immunohistochemistry. The expression of these markers was significantly lower in the Ad-TSG-6 DPSCs than in normal DPSCs (P<0.05). Even without normal DPSCs and TSG-6-RNAi DPSCs induced by TNF-α in vivo, existing vivo results showed that TSG-6 can inhibit the mineralization of DPSCs. The vitro experiment confirmed the results of it in vivo and showed that TSG-6-RNAi DPSCs could have higher mineralization ability in the inflammatory environment. In the future, we will use more methods to prove it in vivo that DPSCs have stronger mineralization capabilities with low expression of TSG-6 in inflammation environment. In addition, TSG-6 gene–modified cells still need to be orthotopically transplanted in animals such as rats, minipigs and rhesus monkeys for further studies.

In conclusion, TSG-6 has inhibitory effect on the BMP-4/Smad signaling pathway and the odonto/osteogenic differentiation of dental pulp stem cells. The results of the present study provided a reference for exploring new treatments for pulp inflammation. In the process of using DPSCs to treat pulpitis, TSG-6 can be modified to increase the differentiation ability of DPSCs in the inflammatory environment, thereby enhancing their repairing ability. In addition, the local use of TSG-6 inhibitors may be conducive to promote the proliferation and differentiation of DPSCs into odontoblast cells in the inflammatory environment, forming the restorative dentin.

## Methods

### Cell culture

The study was approved by the First Affiliated Hospital of Zhengzhou University ethics committee. Human DPSCs were isolated from the third molar tooth of 18-to 25-year-old healthy female patients without caries and inflammation. Then, the tooth pulp was removed under aseptic conditions, using phosphate-buffered saline (PBS; Hyclone, USA), which contained 2% penicillin-streptomycin (Solarbio, China). The removed pulp tissues were washed and cut into tissue blocks of 1 × 1 × 1 mm3. The tissues were digested for 40 min using type I collagenase (Invitrogen, USA) at 37°C. DPSCs were cultured in Dulbecco’s modified Eagle’s medium (DMEM; Hyclone, USA) containing 20% fetal bovine serum (FBS; Hyclone, USA), and the solution was changed every 3 days. After cultivation, the fusion rate was 80%, and the cell passage was conducted. In this study, DPSCs were used to conduct the experiments.

### Osteogenesis and adipogenic induction of DPSCs

Osteogenesis induction: DPSCs were inoculated into 24-well plates, and osteogenic induction solution (10 mmol/L β-sodium glycerol phosphate, Sigma, USA; 10^−8^ mol/L dexamethasone, Sigma; 50 μmmol/L ascorbic acid, Sigma; and 0.01 μmmol/L 1,25-dihydroxyvitamin D3, Solarbio) was added at the fusion rate of 80%. After 21 days, the cells were fixed with 4% paraformaldehyde (Solarbio) and stained with 0.1% alizarin (Solarbio). Then, the mineralized nodule was observed under the light microscope (Olympus, Japan).

Adipogenic induction: The pretreatment of cells was consistent with that in osteogenic induction. The DPSCs were cultured in adipogenic medium (Solarbio, China) for 18 days, fixed with 4% paraformaldehyde, and stained with 0.3% oil red O. The formation of lipid droplets was observed under the optical microscope.

### Flow cytometry analysis

After DPSCs were digested with pancreatin, they were incubated with PE-conjugated antibodies against CD3 (1:100 dilution), CD34 (1:100 dilution), and CD31 (1:100 dilution), and then with FITC-conjugated antibodies against CD90 (1:100 dilution). Moreover, they were incubated with APC-conjugated antibodies against CD73 (1:200 dilution). These procedures were used to determine the expression of cell surface molecules. All antibodies were purchased from BD Biosciences (USA). Flow cytometry was performed using the Biosciences FACSAria III (BD Biosciences).

### Alkaline phosphatase activity assay

A total of 1 × 10^5^ DPSCs were seeded into each well of a 6-well plate. At 70% confluence, DPSCs were cultured in the mineralization-inducing fluid (5% FBS, 10 mmol/L β-sodium glycerol phosphate, Sigma; 10^−7^mol/L dexamethasone, Sigma; 50 ng/mL ascorbic acid, Sigma) containing 0, 10, 50, and 100 ng/mL TNF-α separately for 7 days. Then, DPSCs were harvested, and total proteins were extracted by adding RIPA lysis buffer (Beyotime, China) without phosphatase inhibitor and quantified using a BCA protein quantification kit to ensure that each group was loaded with 15 μg protein. Moreover, an alkaline phosphatase kit (Beyotime) was used to detect the alkaline phosphatase activity according to the manufacturer’s protocols.

### Real-time polymerase chain reaction

DPSCs were cultured in the mineralization-inducing fluid containing 0 and 50 ng/mL TNF-α separately for 3, 5, and 7 days. Moreover, TRIzol (TaKaRa, Japan) was added to extract RNA. Then, reverse transcription (TaKaRa) was performed. The cDNA was amplified using a fluorescent quantitative polymerase chain reaction (PCR) kit (TaKaRa) and an instrument (Applied Biosystems 7500, USA). In this experiment, the expression of GAPDH, TSG-6, BMP-4, DSPP, DMP-1, and ALP was examined. The PCR program was set at 95°C for 30 s; 40 cycles of 95°C for 5 s and 60°C for 34 s; followed by 95°C for 15 s and 60°C for 1 min. All experimental steps were performed according to the instruction on the corresponding kits, and the relative expression levels were calculated using the 2^-ΔΔCT^ method. Table 1 shows the primer sequences.

**Table 1.**
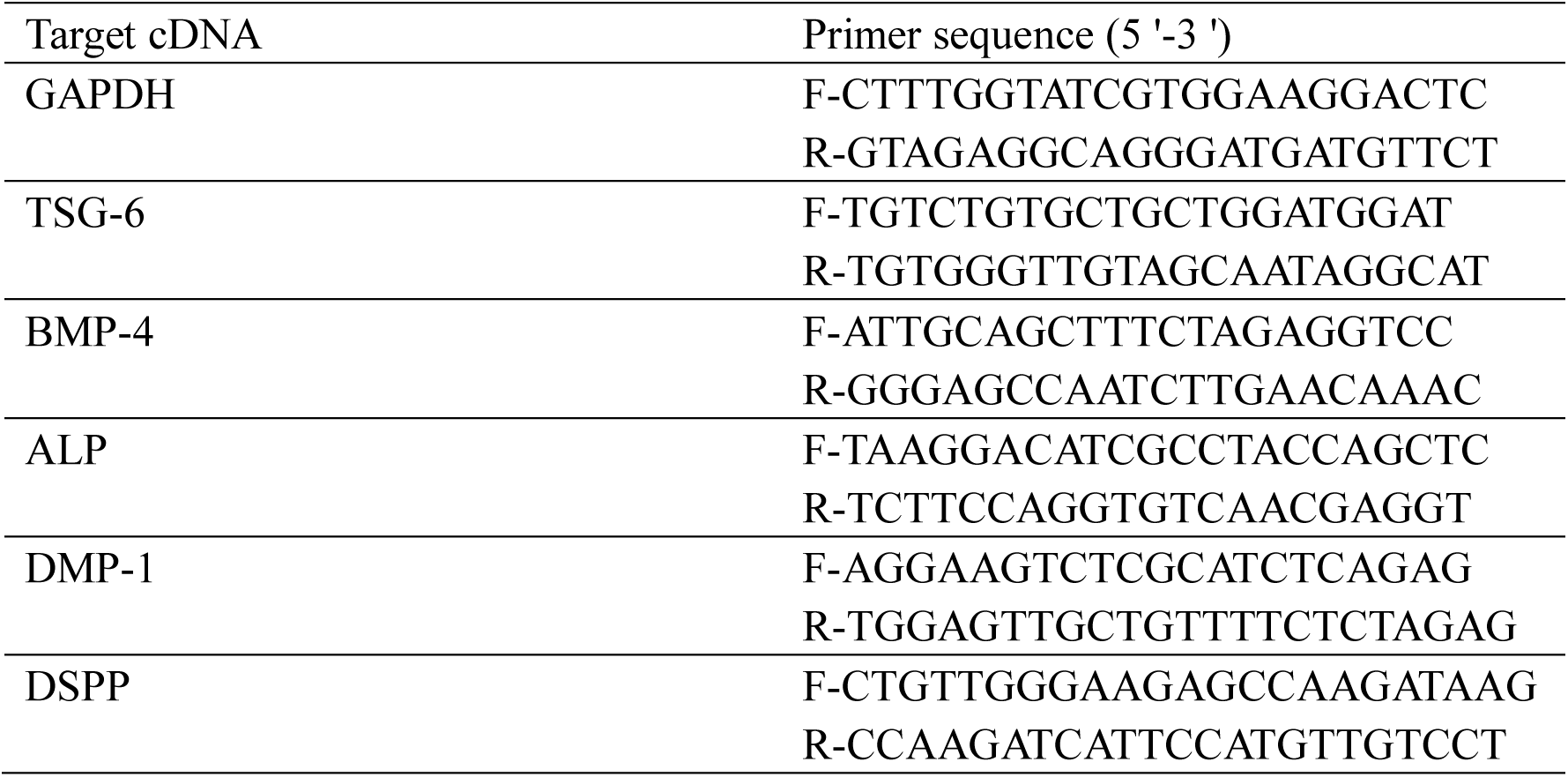
Oligonucleotide primer sequences.

### Transfection

The lentiviruses that overexpress or knockdown the TSG-6 gene were purchased from Shanghai GeneChem (China). Furthermore, DPSCs were inoculated in six-well plates at a density of 1 × 104 cells/well. The lentiviral infection reagent was added after 24 h. The multiplicity of infection (MOI) of 40 could achieve greater than 70% of transfection efficiency. Hence, the MOI was 40 in this study (Supplementary material). After 12 h, the cells were transferred to normal medium culture, and transfection was carried out for 48 h before subsequent experimental steps.

### Western blot analysis

After transfection for 48 h, the TSG-6-overexpressed DPSCs (Ad-TSG-6 DPSCs) were cultured with the mineralization-inducing fluid, and the TSG-6 knockdown DPSCs (TSG-6-RNAi DPSCs) were cultured with the mineralization-inducing fluid containing 50 ng/mL TNF-α. Then, the expression levels of signaling pathway proteins, including TSG-6, BMP-4, and Smad1/5, were determined 48h later. Moreover, after 14 days, the expression levels of odonto/osteogenesis-related proteins, including DMP-1, DSPP, and ALP, were determined. The total proteins were extracted using a protein extraction kit (Dingguo, China). The proteins of each group were separated on a polyacrylamide gel, transferred to a PVDF membrane (Millipore, USA), and blocked with 5% BSA (Solarbio) for 2 h at room temperature. Then, the cells were incubated with primary antibodies, including BMP-4 (1:1000 dilution, Abcam, UK), TSG-6, (1:500 dilution, Abcam), Smad1/5 (1:1000 dilution, CST, USA), DSPP (1:300 dilution, Santa Cruz, USA), DMP-1 (1:300 dilution, Santa Cruz), ALP (1:300 dilution, Santa Cruz), and GAPDH (1:10,000 dilution, Zenbio, China) overnight at 4°C and then with the secondary antibody horseradish peroxidase–conjugated anti-rabbit or anti-mouse IgG antibody (Biyuntian, China) at room temperature for 1 h. The labeled proteins were visualized using a GE Amersham Imager 600 (USA) instrument. Furthermore, ImageJ (National Institutes of Health,USA) was used to analyze the gray value for quantifying results. The protein levels were statistically quantified using the target gene/GAPDH.

### Immunofluorescence

Cell identification: DPSCs were inoculated in 24-well plates and fixed with 4% paraformaldehyde for 30 min. Then, they were permeabilized for 10 min in 0.1% Triton-100 (Solarbio) and washed with PBS four times and then with 10% goat serum. (Solarbio) The samples were blocked for 1 h at room temperature. The cells were incubated with primary antibodies (anti-Vimentin, 1:200 dilution, Abcam; anti-cytokeratin 14, 1:200 dilution, Millipore, USA; anti-STRO-1, 1:100 dilution, Santa Cruz) at 4°C overnight and washed with PBS four times. The incubation with the secondary antibody was performed at room temperature in the dark (cy-3-conjugated goat anti-mouse, 1:500 dilution, Beyotime; FITC-conjugated goat anti-rabbit, 1:500 dilution, Beyotime) for 1h. Finally, 10 µg/mL DAPI (Solarbio) was added to stain the nuclei at room temperature for 10 min.

The transfected and untransfected DPSCs were plated in 24-well plates. The Ad-TSG-6 DPSCs were cultured with the mineralization-inducing fluid, while the TSG-6-RNAi DPSCs were cultured with the mineralization inducer containing 50ng/mL TNF-α for 14 days. The operational steps were the same as for cell identification. Moreover, the primary antibodies were as follows: DMP-1 (1:200 dilution, Santa Cruz) and DSPP (1:200 dilution, Santa Cruz). The secondary antibodies were as follows: cy-3-conjugated goat anti-mouse (1:500 dilution, Beyotime). Cell immunofluorescence was carried out under a laser confocal microscope (LSM800, Zeiss, Germany). Then, the average fluorescence intensity was measured directly using laser confocal microscopy (ZEN2.4 system).

### Subcutaneous transplantation

The animal experiments were divided into four groups to compare the effects of the TSG-6 gene modification of DPSCs on their odonto/osteogenesis differentiation ability. Normal DPSCs were induced with mineralization-inducing solution; Ad-TSG-6 DPSCs were induced with mineralization-inducing solution; normal DPSCs were induced with mineralization-inducing solution containing 50 ng/mL TNF-α; and TSG-6-RNAi DPSCs were induced with mineralization-inducing solution containing 50 ng/mL TNF-α. After culturing the cells for 7 days in the pre-culture medium, they were mixed with PuraMatrix peptide hydrogel (354250, Corning, USA) and injected into the immunodeficient mice at 1 × 10^6^ cells/injection point (6-week-old males, *n* = 12) in the back. The nude mice (Balb/c-nu) were purchased from Beijing Vital River Laboratory Animal Technology Co., Ltd. Six weeks later, all samples were collected from immunodeficient mice, fixed overnight with 4% paraformaldehyde, and then embedded in paraffin. Paraffin sections were prepared and subjected to hematoxylin and eosin, Masson trichrome, and immunohistochemical staining.

The antibodies used for immunohistochemistry included DMP-1 (1:100 dilution, Santa Cruz), DSPP (1:100 dilution, Santa Cruz), COL-II (1:500 dilution, Servicebio, China), OCN (1:500 dilution, Servicebio), and OPN (1:500 dilution, Servicebio). The secondary antibodies were visualized using a DAB chromogenic kit (Zhongshan Golden Bridge Biotechnology, China). Moreover, ImageJ (National Institutes of Health) was used to count the positive signals of immunohistochemical images.

### Statistical analysis

All data were expressed as mean ± standard deviation. Statistical differences were analyzed using the SPSS 21.0 software (SPSS, USA). Student paired *t* test and one-way analysis of variance were used to determine the level of significance. A *P* value less than 0.05 was considered to be statistically significant.

## Acknowledgement

National Natural Science Foundation of China (31670994); Nature science fund of Henan province(182300410340)

